# Unsupervised Machine Learning for Analysis of Coexisting Lipid Phases and Domain Growth in Biological Membranes

**DOI:** 10.1101/527630

**Authors:** Cesar A. López, Velimir V. Vesselinov, Sandrasegaram Gnanakaran, Boian S. Alexandrov

**Affiliations:** Theoretical Division, Los Alamos National Laboratory, Los Alamos, NM 87545, USA; Earth and Environmental Sciences Division, Los Alamos National Laboratory, Los Alamos, NM 87545, USA

**Keywords:** Lipid Phases, Machine Learning, Nonnegative Matrix Factorization

## Abstract

Phase separation in mixed lipid systems has been extensively studied both experimentally and theoretically because of its biological importance. A detailed description of such complex systems undoubtedly requires novel mathematical frameworks that are capable to decompose and categorize the evolution of thousands if not millions of lipids involved in the phenomenon. The interpretation and analysis of Molecular Dynamics (MD) simulations representing temporal and spatial changes in such systems is still a challenging task. Here, we present a new unsupervised machine learning approach based on Nonnegative Matrix Factorization, called NMFk, that successfully extracts physically meaningful features from neighborhood profiles derived from coarse-grained MD simulations of ternary lipid mixture. Our results demonstrate that leveraging NMFk can (a) determine the role of different lipid molecules in phase separation, (b) characterize the formation of nano-domains of lipids, (c) determine the timescales of interest and (d) extract physically meaningful features that uniquely describe the phase separation with broad implications.

## INTRODUCTION

Cell membranes contain mixtures of different lipid types, dynamically arranged, that play a key role in various mechanisms responsible for cell survival ^1^. In the past, membranes were thought to be homogenous systems, however, new data suggests that under different stimulus, the lipids can segregate ^2-3^ into detergent-resistant domains commonly called “rafts”. These domains are highly dynamic, varying in size and composition ^4-5^. Importantly, this lateral partitioning is responsible for activation and functioning of membrane-embedded proteins ^6-7^.

While it is possible to experimentally visualize the structure of segregated lipid domains ^2, 8^, a detailed description of such phases are inherently limited by the resolution of the experimental techniques. At this respect, molecular dynamics (MD) simulations provide a molecular understanding of membrane behavior. In fact, the presence of such lateral rearrangement has been studied extensively using coarse-grained (CG) MD of ternary lipid mixtures ^9^. These CG simulations suggest partial segregation in large lipid systems that mimic the lipid variability of the real cell membranes^10^. MD simulations of such realistic biomolecular systems usually contain millions of particles, even when simplified models are used. When the purpose of such simulations is to gain biological or physical insights, it is challenging to identify patterns in the behavior encoded in the motion of thousands of molecules, so developing analytic tools for extracting functionally relevant features from MD generated trajectories is of great importance.

Currently, machine learning (ML) methods have shown a lot of promise in many fields^11^, and recently some of them have been applied for analysis and detection of classical and quantum phase transition data generated by simulations. Both supervised and unsupervised ML approaches have been used for this purpose ^12-19^, but most of these pioneer studies used small Ising-like systems for their investigations. ML has previously been coupled with MD simulations of biomolecular systems in a limited context. ML techniques were reported to predict free-energy differences when trained with MD simulation data ^20^, and unsupervised approaches such as PCA ^21^ as well as other techniques ^22-23^ have been used to reduce the dimensionality of MD generated data ^24-28^.

In general, the unsupervised ML methods learn relationships between elements in uncategorized data and classify the data without human’s help but by revealing its internal structure and latent (i.e., not directly observable) features hidden in the data. The unsupervised methods include clustering ^29^, classical neural networks ^30^, and the more contemporary blind source separation (BSS) techniques ^31^. BSS is based on factorization which is one of the most powerful tools for extraction of latent features ^32^. BSS include principle component analysis (PCA) ^33^, singular value decomposition (SVD) ^34^, and more advanced methods, such as independent component analysis, ICA ^35^ and nonnegative matrix factorization, NMF ^36^.

A limitation shared by PCA, SVD and ICA is the difficulty to relate the extracted latent factors to physically interpretable quantities; NMF overcomes this limitation because the nonnegativity of the extracted latent factors leads to a collection of strictly additive components that are sparse and parts of the data and hence are amenable to a simple and meaningful interpretation without introducing prior assumptions ^37^. NMF decomposes given data matrix, *X*_LN_, into two matrices, W_LK_, and H_KM_, such that X_LN_ ≈ W_LK_ H_KM_. The factor matrices, WLK and H KM, are both nonnegative and have one small dimension *K* that represents the number of the latent features in the data (Fig. 1). A mathematically rigorous formalism is given in the later sections. NMF ability to identify easy interpretable latent features enables discoveries of new causal structures and unknown mechanisms hidden in the data as discussed in the literature ^38^. Surprisingly, the implementation of NMF for analysis of MD simulations at the interface of physics and biology has been lacking.

**Figure 1.**
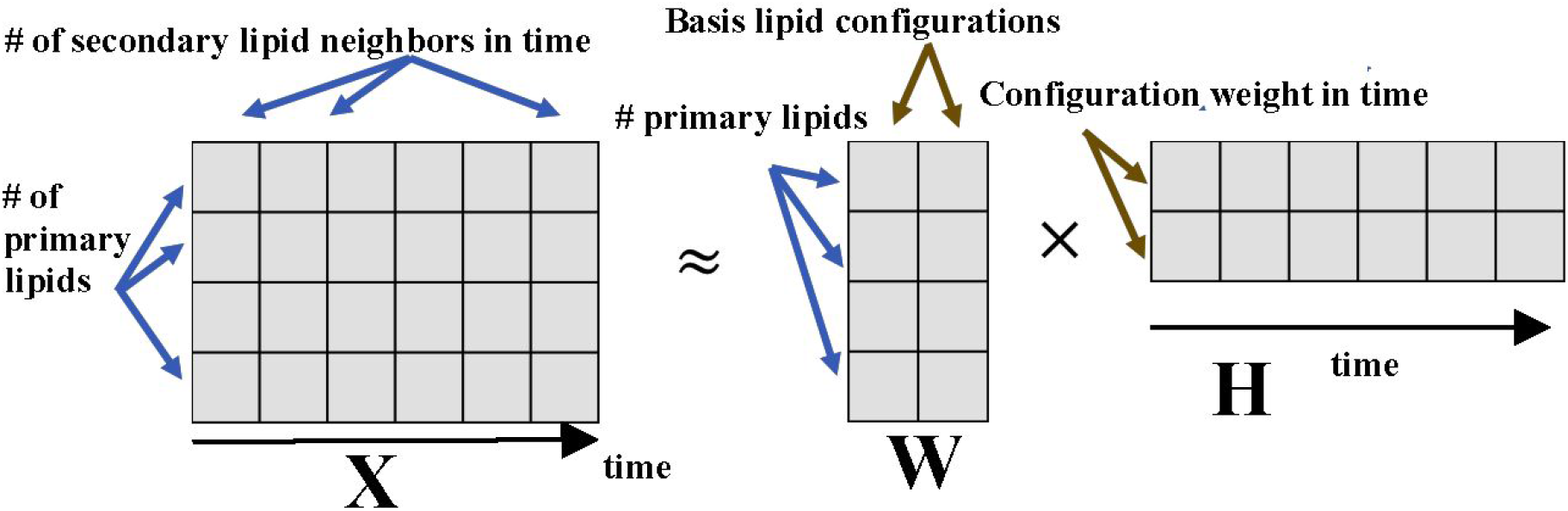
Illustration of a Nonnegative Matrix Factorization. The nonnegative matrix **X** is decomposed to the product of a nonnegative matrix **W**, containing **K**=2 basis lipid configurations, and nonnegative matrix **H**, containing the contributions of these two configurations in different time points.

Here, we present a new unsupervised machine learning algorithm based on NMF combined with custom k-means clustering, called NMFk, capable of analyzing phase separation in a system of mixed lipids directly from the pre-processed trajectories derived by MD simulations. We use CG MD simulations of a physical system that comprises a 3- component lipid mixture, commonly accepted to mimic the behavior of a cellular plasma membrane ^5^. We show that NMFk, applied to a pre-selected data from these simulations is able to, (a) determine the molecules that play different roles in phase separation, (b) characterize the formation of lipid nano-domains, (c) reveal the timescales of interest, and (d) extract physically meaningful features that characterize the phase separation.

## RESULTS

### Generation of lipid mixture data sets using coarse-grained MD simulations

For MD simulations of membranes and membrane-based biological systems, the Martini coarse-grained (CG) force field ^39^, considerably reduces the computational cost of calculations by nearly three orders of magnitude compared with similar MD simulations using fully atomistic force fields ^40^. Particularly, the CG approach can capture relevant dynamics and fluctuations of larger membrane patches, which are prohibitive with atomistic simulations. Such access to larger spatial and temporal scales enables direct comparisons with experimental measurements ^41^. Pioneering computational studies with the Martini CG force field allowed the characterization of not only lipid segregation and lipid phases, but also the relative partitioning of membrane proteins between these phases ^9, 42-44^. More recently, Martini has been used in simulations of membranes with lipid compositions of comparable complexity to those found in specific tissues of living cells ^45-46^.

Regardless of the extensive use of the Martini force field, the building-block principle of Martini along with the 4:1 atoms to bead mapping unavoidably reduce the accuracy due to loss of detailed description of specific molecular chemical properties. Thus, despite the fact that many of the current Martini lipid parameters are sufficient to guide accurate membrane simulations ^39, 47-48^, global lipid properties are compromised ^49^. Consistent with previously published work ^42, 50-51^, the standard Martini V2.2 parameters for DPPC, DOPC, and CHOL do not phase separate at 298 K, or even at 290 K in the physiologically relevant L_d_/L_o_ coexistence region of the phase diagram. Recently, we incorporated changes in the lipid Martini force field ^52^, greatly improving the lipid segregation in line with current experimental phase diagrams ^8, 50^.

We use the above two versions of the Martini force field, one lipid parameter set that phase separates and the other that does not, to generate two data sets for the analysis using NMFk. The first set considers the data from the coarse-grained MD simulations of the current Martini force field (called “Standard”). As mentioned above, the current Martini V2.2 lipid parameters for DPPC and DOPC are not able to properly segregate the DPPC:DOPC:CHOL mixture. It serves as the prototype for non-phase separating homogeneous lipid mixture. The second set considers the refined version of the Martini (called “Updated”), which has been optimized to reproduce the experimental phase separation and domain formation for this ternary system ^52^. It serves as the prototype for phase separating lipid mixture. Using these two versions of force fields, we carried out 20 μs long CG MD simulations. The expectation is that we should able to detect and analyze features associated with phase separation in one case but not in the other case.

### Conventional Analysis of Domains in MD Simulations of Ternary lipid Mixtures

The formation of lipid domains has been heavily studied both experimentally and computationally ^2-4, 8-10, 53-55^ Computational observation of explicit lipid segregation at nearly atomic detail dates back to almost 10 years ago ^9^ Analysis of such processes involved direct visualization of cholesterol-rich/-poor domains, as well as physical quantification of the area per lipid, cholesterol content, radial distribution functions and membrane thickness mismatch ^9^. These analyses brought enough details that membrane domains could be discriminated at the nanoscale. A more sophisticated approach can be cited from the work of Baoukina et al.^51^, where a Voronoi tessellation methodology was applied in order to delineate the boundaries between ordered/disordered domains in monolayers. This approach can be also directly combined with automated predictive tools like Markovian based methodologies ^54^. Regardless of these published methodologies, the analysis and prediction of such fluctuating membrane domains become inaccessible when the complexity of lipid content increases with larger size membrane patches (e.g. plasma membrane).

A typical process of molecular lipid phase separation is depicted in **Fig. 2**. Initially, the lipid components are randomized, mimicking the homogeneity at a high-temperature (**Fig. 2A**). Subsequent fast quenching of the mixture to 290 K, well below the melting temperature of the fully saturated DPPC lipid, leads to the rapid formation of nanoscale domains on a submicrosecond time scale with the Updated Martini force field (**Fig. 2B top panel**). These nano-domains are eventually formed over the entire surface of the membrane, but in different regions. After 0.5 microseconds, the nano-domains start to interconnect, leading to the formation of larger cholesterol-rich regions. In agreement with the general raft hypothesis ^53^ and previous computational studies ^9^, the ‘‘ordered’’ nano-domains contain most of the saturated lipids together with cholesterol forming a Liquid ordered (L_o_) domain, whereas the ‘‘disordered’’ nano-domain is mainly composed of the polyunsaturated DOPC lipid segregated in a Liquid disordered (L_d_) region. Contrary to the Updated force field, the lipid mixture based on the Standard Martini force field does not show any tendency for phase separation or domain formation (**Fig. 2B, bottom** panel), in agreement with previously published data ^52^.

**Figure 2.**
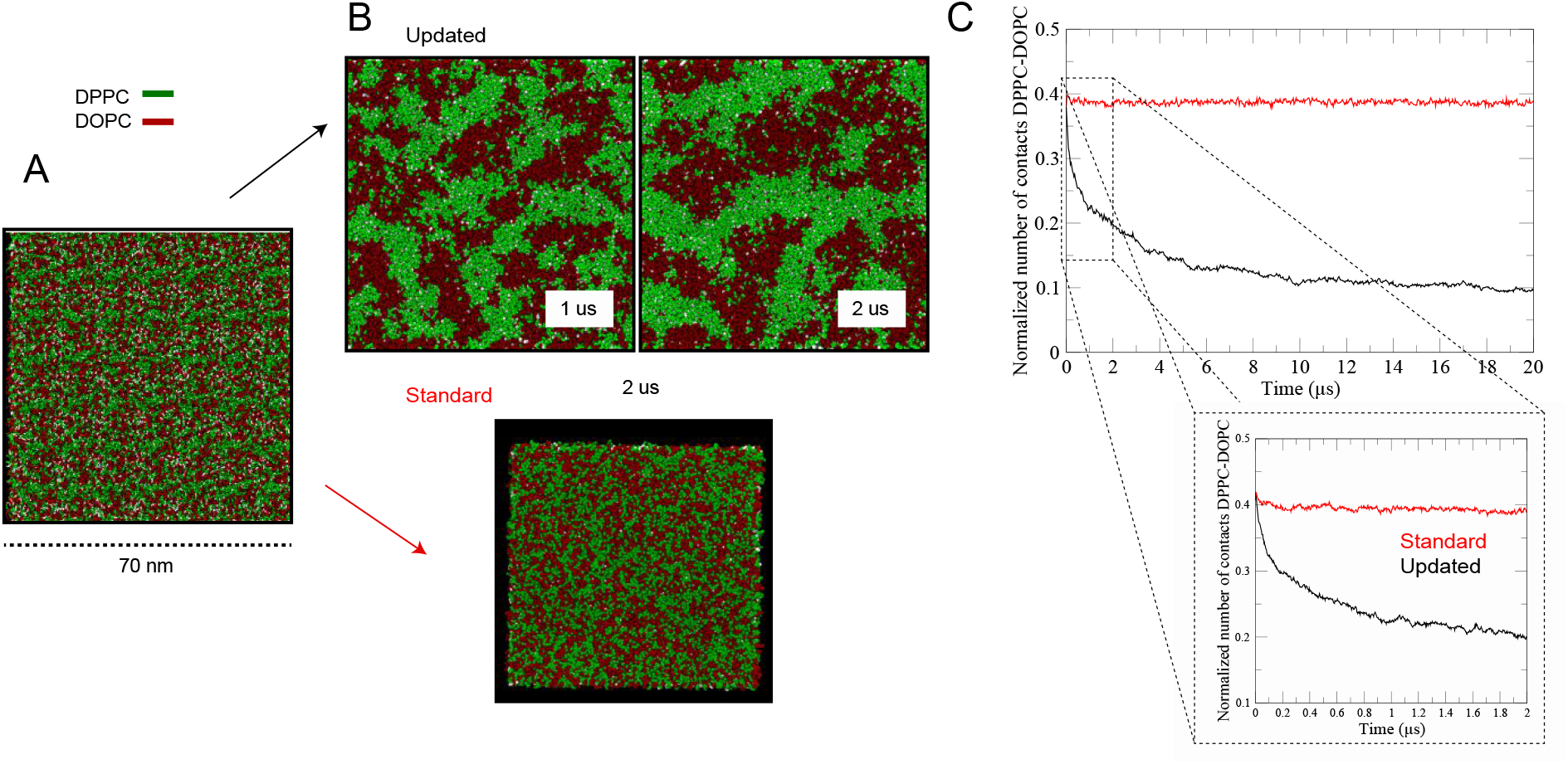
Corse-grained simulations of ternary lipid mixture. **(A)** Initial system setup for the CG simulations. Lipids were initially randomly placed within the XY plane. Saturated lipid, DPPC, is colored in green while unsaturated lipid, DOPC, is colored in red. Cholesterol is colored in white. Water is not shown for clarity in depiction. **(B**) Lipid phase separation within 2 us simulation. The updated CG force field shows clear phase separation into L_o_ (green) and L_d_ (red). Contrary, the standard Martini lipid force field does not show preferential segregation in cholesterol rich domains. **(C)** Time evolution of the normalized number of contacts between saturated and unsaturated lipids, showing poor separation with the standard Martini force field.

Following the conventional approaches to quantify the segregation tendency, we compute the normalized total contacts between DPPC and DOPC as a function of simulation time (**Fig. 2C)**. Initially, these contacts are featured by larger values, meaning that these two lipid types are indeed in close contact, highlighting the initial homogeneous lipid mixing of the system. However, with the Updated Martini, lipid segregation leads to a decrease in the total number of contacts between DPPC and DOPC (**Fig. 2C black line**). Whereas, the contacts remain unchanged in the simulations with the standard Martini (**Fig. 2C red line**). In the case of Updated Martini, contacts decay during the simulations and begin to plateau with increasing simulation time. We should note here that the proper convergence to a stationary state may not be achievable within the simulation timescales considered here, as already published ^54^, while significant transition towards segregation occurs within the first 2 us (**Fig 2C insert**). We consider this time regime suitable for NMFk analysis to extract latent features associated with the phase separation.

### NMFk implementation for analysis of ternary lipid mixture simulations

To perform the NMFk analysis, we first define the primary and secondary lipid types and then calculate the neighbor matrix at a given time ***t***, ***X_L_***(***t***), in terms of the number of secondary lipid type surrounding the primary lipid type. For this study, we considered DPPC and DOPC as the primary and secondary lipid types, respectively, although, there is no restriction of other combinations for primary and secondary lipid types. We compute the number of closest DOPC neighbors, ***X_L_***(***t***), around every DPPC lipid, *L*=(*L*_1_,*L*_2_, …, *L_M_*), within the distance corresponding to the second peak of the DOPC-DPPC radial distribution function (**Fig. 3 blue dashed line**). The number of neighbors within this distance serves as the order parameter for the lipid phase separation. This neighbor data is recorded at *N* consecutive simulation time points, *t*=(*t*_1_,*t*_2_, …, *t*_*N*_),> corresponding to the time evolution of the system up to first 2 us. The matrix ***X_L_***(***t***) contains *N* arrangements at *N* time points, presented by the columns of ***X***, that form the matrix of the neighbors, ***X***_𝑳_ ***t***; ***X*** ∈ *M*_*MN*_(𝐼_+_), where *M* is the number of the rows, and each row represents the time evolution of the closest neighbors of one specific lipid of the primary type. The *I*_+_ denotes the set of nonnegative integer numbers. Next, NMFk decomposes the neighbor matrix ***X_L_ t*** as a product of two matrices: ***X_L_***(***t***) ≈ ***W_LK_*** ∗ 𝑯_***K***_(***t***), where the columns of ***W_LK_*** are the ***K*** basis lipid configurations describing the state of the lipid system and 𝑯_***K***_(***t***) contains the contributions of each one of these configurations at time ***t*** (see Fig. 1 where K=2).

**Figure 3.**
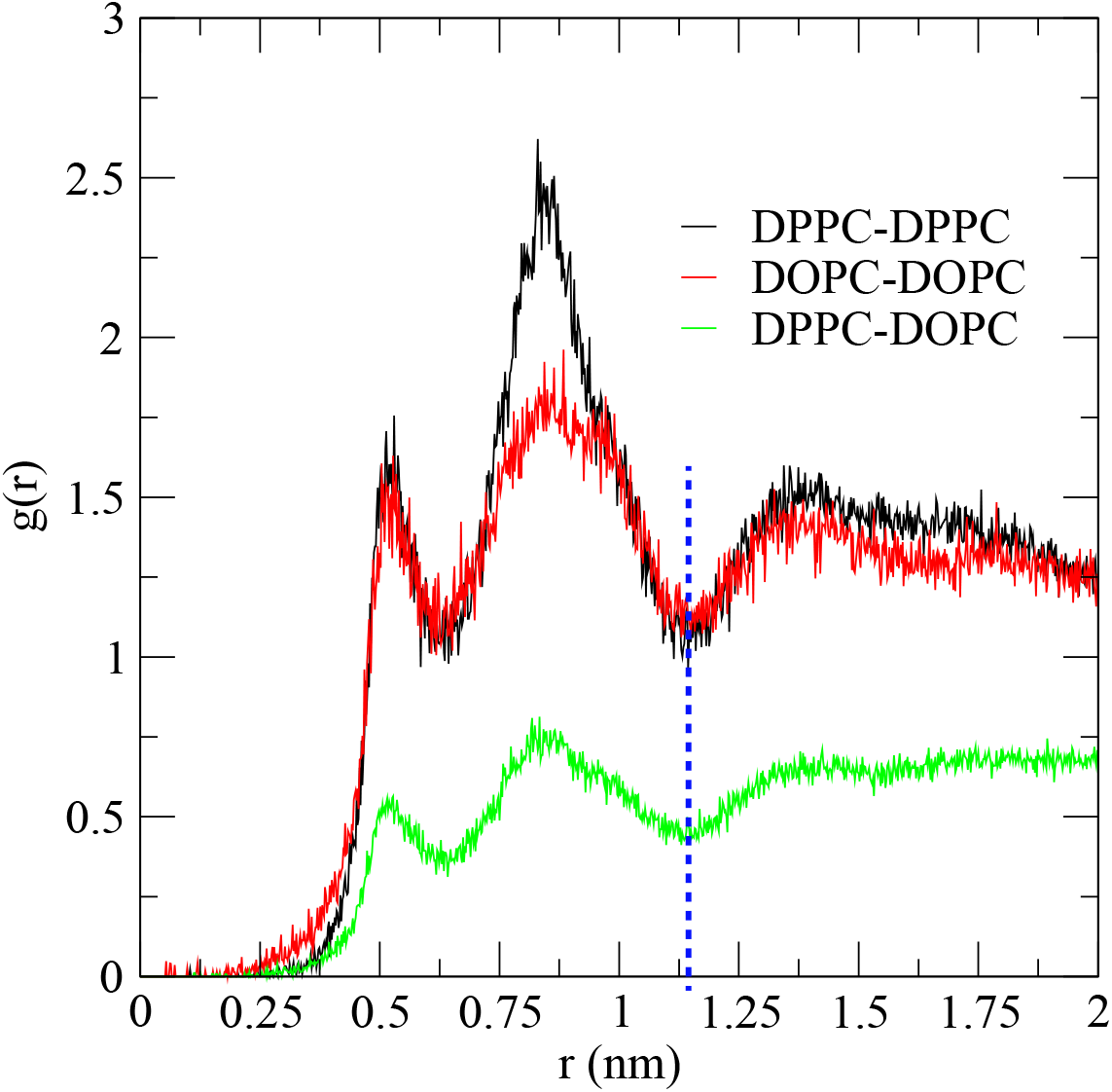
Lateral radial distribution function for the different lipid combinations. RDF was computing considering the center of mass of the molecules. The dashed blue line indicates the chosen cut-off distance for profiling neighbor list needed for the NMFk matrix construction.

The neighbor matrix, ***X_L_***(***t***), contains relevant properties of the ternary lipid mixture. At the beginning of the simulations, the lipid mixture is homogeneous and there are no distinct phases, i.e., there is no specific structure in ***X_L_***(***t***). Hence, ***X_L_***(***t***) contains uniformly distributed integer numbers that do not have any distinct features. With the Updated Martini force field, by 2 us, the primary lipid type, DPPC, segregates into the L_o_ region, while the secondary lipid type, DOPC, segregates into the L_d_ domain. In this case, if a concrete lipid of the primary type is located deep in a L_o_ domain, the probability to have a lipid-neighbor of secondary type is small (close to zero). In contrast, if this primary lipid is situated outside of any domain, the probability to have a neighbor of the secondary type is much higher. Therefore, we expect the neighbor matrix to contain a structure that tracks this phase separation and the subsequent analysis by NMFk to extract the hidden features describing the structure and the phase separation.

The NMFk analysis, as we will see next, demonstrates exactly that: When phase separation begins at time, ***t*=*t***_***s***_, for each one of the snapshots recorded at, ***t* > *t*_*s*_**, and for each of the 𝑴 lipids of the primary type, the values of the matrix ***X***_𝑳_(***t*** > ***t***_***s***_) can be represented as a linear mixing of ***K*** basis lipid configurations presenting the probability of the given lipid of the primary type, 𝐿_𝑖_, to have a closest neighbor-lipid of secondary type. NMFk decomposes the matrix ***X_L_***(***t***), to a nonnegative probability matrix, 𝑾, 𝑾 ∈ *M*_*M*𝐾_(ℛ_+_), corresponding to these ***K*** basis lipid configurations, blended by the weights, presented by the elements of a nonnegative matrix, 𝑯, 𝑯 ∈ *M*_𝐾*N* − 𝑠_(ℛ_+_) that reflects how these configurations are active and mix in time. Thus, for a given arrangement of the primary lipids, ***X***_𝑳_, at a time point ***t* > *t_s_***, we have,

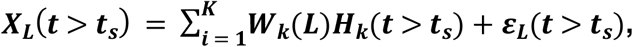

where 𝜺 ∈ *M*_*MN* − 𝑠_(ℛ_+_) denotes the presence of a noise or unbiased error of decomposition. Before the time point ***t*** = ***t_s_***, there is no trace of a phase separation and the pattern of the number of the closest neighbors is stochastic. Therefore, there is no clear features that could be recognized by NMFk.

The reconstruction of ***X***_𝑳_(***t*** > ***t_s_***), via the two factor matrices, 𝑾_𝒌_(𝑳) and 𝑯_𝒌_(***t*** > ***t_s_***), serves as a measure for the significance and quality of the extracted latent features. For the case of Updated Martini, **Table 1** presents the Pearson correlation coefficient between the reconstructed lipid arrangements at each time point ***t*** (obtained by NMFk) and the original arrangements at the same time point (i.e., the corresponding column of the neighbor matrix ***X_L_***(***t***)). The ***K*** unique basis lipid configurations (encoded in the probability matrix 𝑾), reproduced the 𝑵 − 𝒔 lipid arrangements forming the matrix ***X***_𝑴𝑵 − 𝒔_. With the standard Martini, the NMFk was not able to provide a set of lipid configurations that can reconstruct the simulations accurately. Indeed, and as shown in **Table 1**, the NMFk analysis did not reproduce accurately the simulation data.

**Table 1.** The quality of reconstruction of X, estimated by the mean Pearson correlation between the columns of X and columns of W*H.

| Coarse-grained simulations of ternary lipid mixture | Time [us] | Mean Correlation Coefficient | Standard Deviation |
| --- | --- | --- | --- |
| Standard Martini | 0.45 - 2 | 0.54 | 0.040 |
| Updated Martini | 0.45 - 2 | 0.90 | 0.020 |
|  | 16 - 18 | 0.96 | 0.004 |

### NMFk derived features associated with phase separation of ternary lipid Mixture

Here, we describe how the basis lipid configurations extracted by NMFk describe physical properties of the phase separation, such as, time profile of which lipid molecules belongs to which phases, the formation of nano-domain, and the spatial and temporal profiles of nucleation and phase separation.

A typical outcome of NMFk analysis is presented on **Fig. 4**. According to our Silhouette-Reconstruction criteria (see Methods Section), NMFk determines that the optimal number of basis lipid configurations is 20 (**Fig. 4A**). The columns 𝑯_𝒊_ of the matrix 𝑯, each of which encodes the weights, that is, the participation of a basis lipid configuration in time, are presented on (**Fig. 4B**). It is clear that for each 𝑯_𝒊_ there is a well-defined time interval containing a number of consecutive frames where the corresponding basis configuration 𝑾_𝒊_ is active. From **Table 1**, it can be seen that after 0.45 μs, NMFk reproduces the neighbor matrix ***X_L_***(***t***) very well: the cross-correlation between the neighbor matrix and the reconstructed matrix, for each time frame, is above 0.95. The much lower cross-correlation for reconstruction of the matrix ***X_L_***(***t***) at early times suggests a prevalence of homogeneous lipid mixture in the first 0.45 μs. The 15 significant basis lipid configurations that are active at consecutive time intervals after the first 0.45 μs (#6; #10; #12; #15; #17; #19; #20; #18; #16; #14; #13; #11; #7; #2; #1) represent the evolution of the nucleation and phase separation.

**Figure 4.**
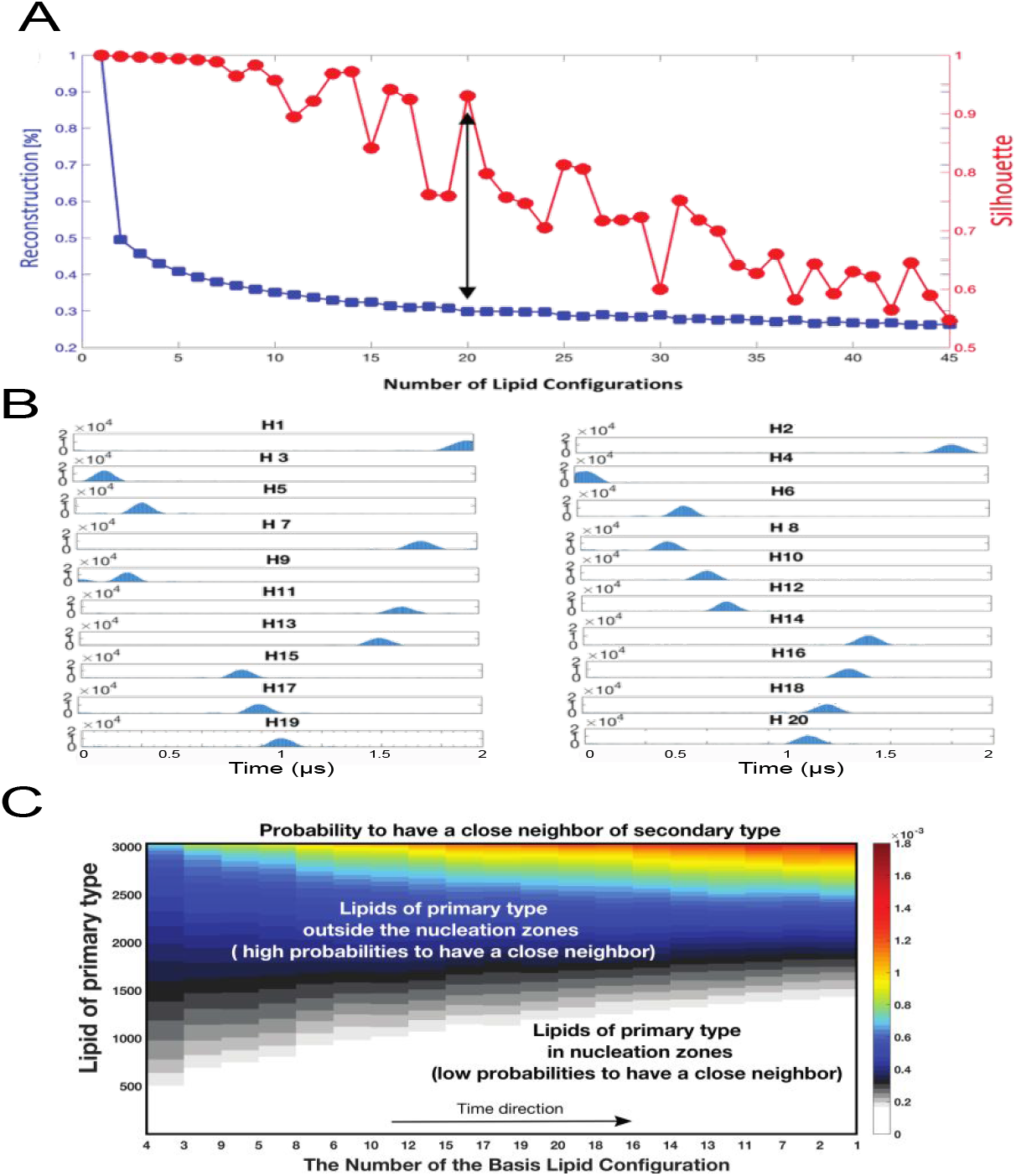
Outcome from NMFk analysis. **(A)** The Silhouette-Reconstruction criterium (see the Methods Section). On the x-axis is denoted the number of basis lipid configurations and on the y-axis-the average Silhouettes (right y-axis, red marking) as well as the Reconstruction (left y-axis, the blue marking). The double arrow denotes NMFk estimates of the number of basis lipid configurations. **(B)** Presentation of the columns 𝑯_𝒊_ of the matrix 𝑯 that encodes the weights of the basis lipid configurations at given time frame. **(C)** A heatmap presenting the basis lipid configurations ordered along x-axis according to the time interval they occurred from early time to late as gathered from the corresponding weight-columns, 𝑯_𝒊_. The color gradient from white to blue to red represents the increase in probability for a specific primary lipid to have a secondary lipid neighbor within each basis lipid configurations. The increase in white and gray colors with time is indicative of the evolution of the phase separation.

Each one of the 15 significant basis lipid configurations, extracted by NMFk, contains a set of probabilities for the lipids of a primary type to have a close neighbor-lipid of a secondary type. Each basis lipid configuration (i.e., the specific set of probabilities) is active at a given time interval defined by the weight of this basis configuration at the corresponding column of the matrix 𝑯. In each basis lipid configuration, we expect to have at least two groups of probabilities: (a) the probabilities of primary lipids situated in the nucleation domains that have relatively small number of neighbor-lipid of secondary type, and therefore these probabilities, on average, approach zero; and (b) the probabilities for primary lipids outside of any nucleation domain whose number of neighbor-lipid of secondary type is much higher. There are always transition of the primary type of lipids in and out of the nucleation domains or at the interface of the nucleation domains: at a certain moment, a given primary lipid could be situated outside a nucleation domain formed by primary lipids, but after a while it could reach interfacial region and eventually get absorbed into the nucleation domain. Alternatively, primary lipids inside the nucleation domain or at the interfacial region may venture out into the liquid disordered region enriched of secondary lipids. These exchanges continuously alter the probability of a given lipid of the primary type to have neighbor-lipid of secondary type, as the phase separation proceeds which results in different basis lipid configurations at different time points as the system goes to a phase separation. At long timescale, when the phase separation has reached equilibrium, basis lipid configurations capture the exchanges of the primary lipids governed by the stochasticity and diffusion.

To better characterize the above observations related to the extracted basis lipid configurations, we applied k-means clustering on each of the 20 basis lipid configurations. We combined the k-means clustering with Silhouette statistics to determine the most probable number of clusters, i.e., the most probable number of groups of probabilities in each basis lipid configuration. Specifically, we calculate consecutively the Silhouettes of the resulting clusters, changing the number of clusters from 1 to 30 to determine the most probable groups of primary (DPPC) lipids with similar probabilities to have a neighbor secondary (DOPC) lipid-neighbor. **Fig. 4C** shows the basis lipid configurations ordered according to the sequence of the frames corresponding to consecutive time intervals determined by their respective weights. The k-means clustering procedure determined that each of the extracted 20 lipid basis configurations can be separated to two clusters: The first cluster contains the primary lipids with a low average probability to have a DOPC lipid-neighbor and the second cluster contains the primary lipids with more than four times higher average probability to have a DOPC lipid-neighbor. Further, we colored differently the lipids in each of the 20 lipid basis configurations, with two clusters each, at the time intervals where the respective lipid basis configuration is active. The color gradient in **Fig. 4C** captures these two groups of primary lipids: white to grey the primary lipids within the nucleation domains, and blue to red-the primary lipids at the interface or outside of any nucleation domain.

Next, we use the MD simulations trajectories to visualize and rationalize the two primary lipid groups as extracted by the clustering of basis lipid configurations derived by NMFk. We considered lipid basis configurations, #19 and #1 corresponding to 1us and 2us time points, respectively. All lipids contributing to those lipid basis configurations are mapped into the trajectories at the corresponding time points in **Fig. 5**. At 2 us, approximately 70% of primary lipids (DPPC, colored in blue) are situated in large nucleation domains which are well packed and condensed by the high cholesterol concentration, leading to L_d_ phase. These primary lipids are predominantly shielded from the secondary (DOPC, colored red) lipids which themselves are localized to form the L_o_ phase. These primary lipids correspond to the first cluster identified by NMFk and are localized in nucleation domains.

**Figure 5.**
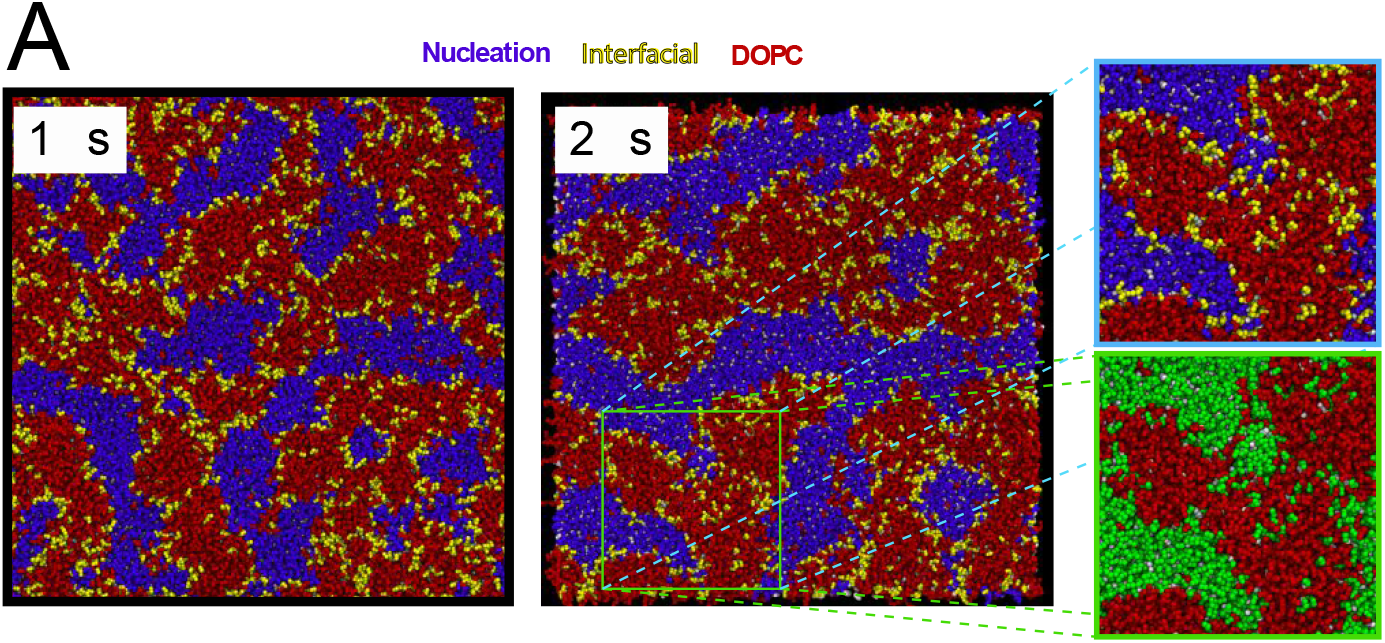
Visual inspection of simulations trajectories for evaluation of NMFk categorization primary lipids according to their localization. Primary (DPPC) lipids are colored according to NMFk output as purple (in nucleation domains) and yellow (out of nucleation domains) in MD configurations corresponding to two time points. Secondary (DOPC) lipids are colored in red and cholesterol in white. Insets highlight a particular region where lipids can be differentiated as nucleating (purple) and boundary (yellow) lipids. As a comparison, the same region is represented using the conventional way of addressing the distinction between saturated (green), unsaturated (red) lipids, similar to Fig. 1.

On the other hand, at 2 us, approximately 30% of the primary lipids (DPPC, colored yellow) in the same lipid basis configuration #1 are located near the interfacial regions or deep into the L_d_ regions formed by secondary lipid types. Unlike the previous set of primary lipids, these lipids are in direct contact with the secondary lipids. The **Fig. 5 insert** shows how NMFk is able to distinguish these lipids from all DPPC lipids available. They correspond to the second cluster identified by NMFk. Thus, each given lipid basis configuration contains physical and easily interpretable features that enable us to make a distinction on primary lipids depending on their location and their contribution to the lipid segregation.

The strength of NMFk analysis is the ability to extract lipid basis configurations as a function of time. These configurations enable us to determine the time dependence of the latent features connected to the kinetics of phase separation without carrying out tedious multiple analyses. A conventional analysis that seeks to probe the temporal profile would have considered the normalized number of contacts among DPPC lipids and between DPPC and DOPC to capture the nucleation process (**Fig. 2C**). NMFk decomposes the nucleation process in two components, as shown in **Fig. 6**, where the distinction is made on the temporal profiles of primary lipids from the nucleation domains from those that are still outside the nucleation domains. In **Fig. 6A**, the purple bars represent the total number of primary lipids in nucleation domains at consecutive time intervals (ordered on the x-axis), while the primary lipids outside of any nucleation domain are presented by yellow bars. At early time, rapid growth of domains is seen which is directly correlated to the increase of nucleation of primary lipids. After this rapid growth, a steadier behavior is observed which continues till the end of the simulation. In **Fig. 6B** we represent the normalized number of contacts with the secondary lipids for the primary lipids as identified from NMFk. At early times, these contacts are high due to random encounters between lipids. This behavior begins to change around 0.6 us and then the contacts between primary and secondary lipids are indicative of steady growth of the number of primary lipids in the nucleation domains and a decrease of the total number of lipids in the non-nucleating regions as the lipid mixture system phase separates.

**Figure 6.**
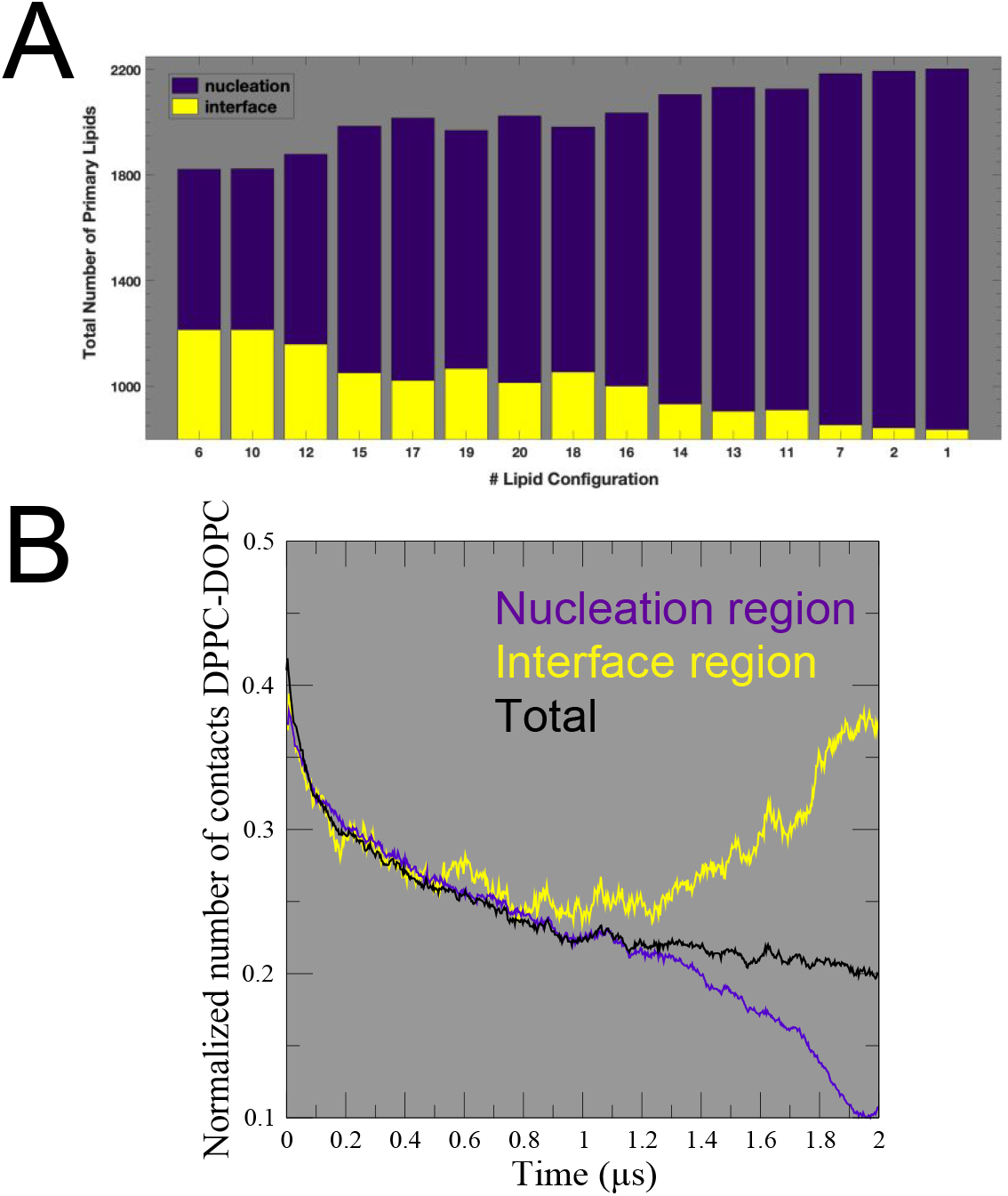
Time dependent variation of the total number of nucleating primary (DPPC) lipids. **(A)** The purple bars represent the total number of primary lipids in nucleation domains, as concluded from their membership in different clusters of the corresponding basis lipid configuration, while the primary lipids outside of any nucleation domain are presented by the yellow bars. The labels on x-axis (the numbers) correspond to consecutive (in time) processes extracted from 0.45 us to 2 us simulation time. **(B)** The same distinction of nucleating primary lipids as in **(A)** obtained using the normalized number of contacts of the lipids identified by NMFk. The same color scheme is used. Black line corresponds to the normalized number of contacts using the total fraction of saturated lipids (i.e., without making the distinction within the nucleating primary lipids).

Furthermore, NMFk analysis extracts the temporal evolution of the primary lipids’ *membership:* in or out of a nucleation domain. Specifically, the membership is defined by identifying primary lipids that join a nucleation domain and remain in that domain till the phase separation. We ordered the 15 significant basis lipid configurations extracted by NMFk according to the time intervals when the specific configurations are active, **Fig. 7**. We identify the primary lipids that participate in the nucleation by keeping track of the lipids in basis lipid configurations with low probabilities to have a secondary lipid-neighbor. In **Fig. 7A**, we present the *concrete* primary lipids that *remain* in the same cluster with small (purple color) or high (yellow color) probability to have a secondary lipid-neighbor at consecutive time intervals, which represents the evolution of nucleation. The inset in **Fig. 7A** demonstrates the system at much later time (∼ 20 microseconds) after the initial nucleation when the phase separation has reached equilibrium and the primary lipids that are located in nucleation domains mostly preserve their membership in time. Importantly, this evolution of the primary lipids can be easily mapped at their spatial coordinates. In **Fig. 7B**, the patterns extracted by NMFk are mapped via MD trajectories to their spatial coordinates (at each time interval), to visualize the evolution of the different groups of primary lipids based on their membership: in nucleation domains (purple color) or outside those domains (yellow color). This again highlights the power of the NMFk to extract physical properties and details that can be easily visually tracked as the system undergoes localization and lipid segregation.

**Figure 7.**
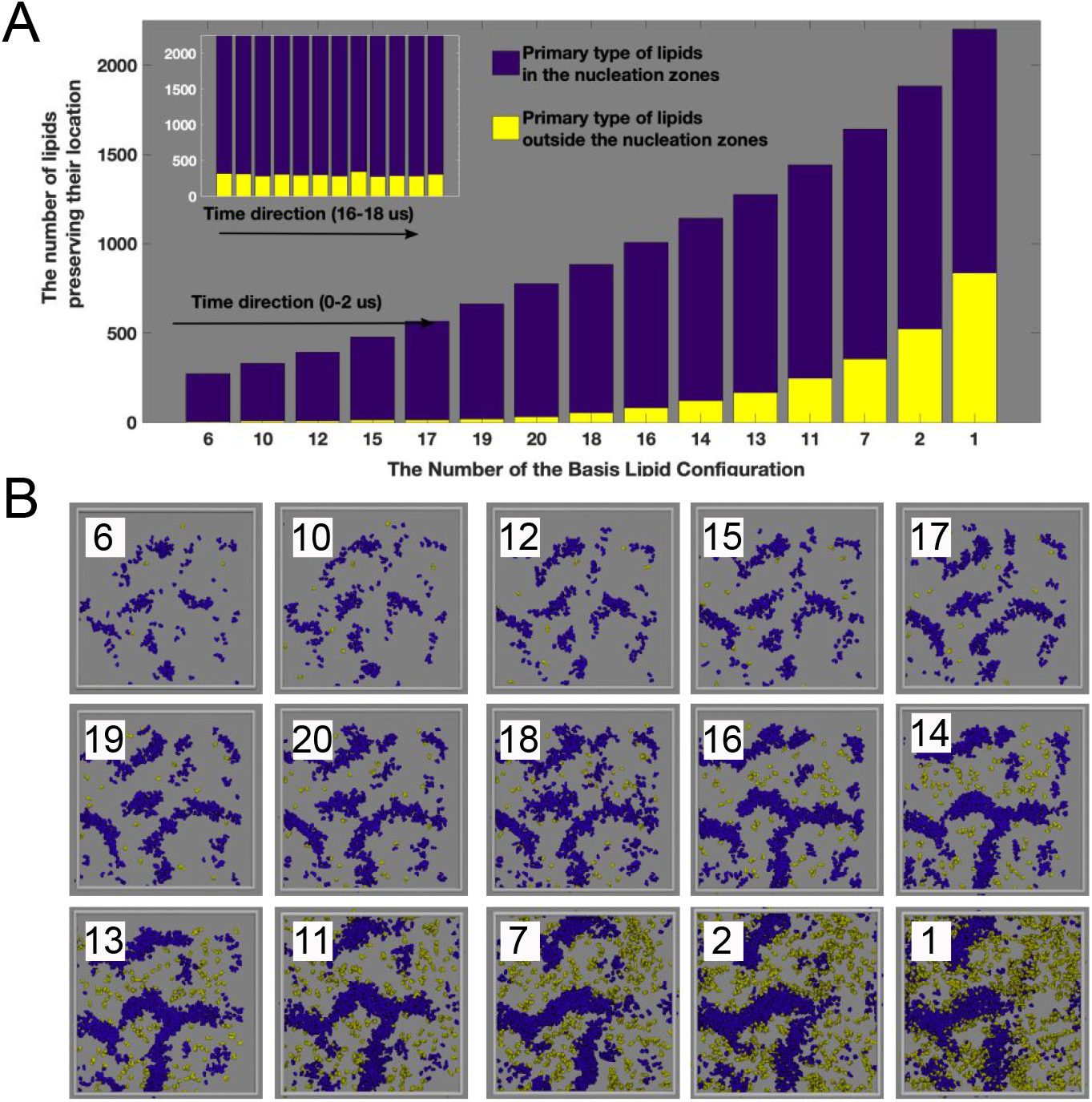
Temporal description of membership profile of primary lipids as captured by NMFk analysis. **(A)** The clear growth of nucleation domains with time, formed by primary lipids, is extracted from the clusters containing primary lipids with low probabilities to have lipid-neighbors of secondary type (refer Fig. 4C). The number of primary lipids that belong to nucleation domains are colored in purple whereas those appear at the interface (edges of the nucleation domain) or outside the nucleation domain are colored in yellow. The exponential growth of the number of the primary lipids that continue to be in the same cluster demonstrated the evolution of the steady state of the phase separation. The inset demonstrates the same process but at much later time (∼ 20 microseconds) when the phase separation is in equilibrium. Here, although the minor changes still exist, primary lipid membership numbers have stabilized. **(B)** Spatial visualization of the primary lipid memberships extracted by NMFk analysis (between 0.45 us and 2 us). These primary lipids were tracked using MD trajectories and rendered with the same color scheme as in **(A)** to distinguish the primary lipids in and out of nucleation domains. Other lipids are not rendered (silver background). The MD simulation box is represented as solid gray lines.

## DISCUSSION

We introduce an unsupervised machine learning algorithm based on the nonnegative matrix factorization combined with custom clustering, called NMFk, for analysis of MD simulations. Specifically, we implement that algorithm here to detect and describe the lateral lipid segregation in a simplistic lipid “raft” model composed of a well-characterized ternary lipid mixture. Based on this study, we believe that the NMFk formalism can be also implemented for extracting relevant features from a more complex biological membrane.

The DOPC:DPPC:CHOL ternary lipid mixture considered here can exist as a homogeneously mixed mixture or exhibit two distinct phases of L_o_ and L_d_ depending on temperature. NMFk is utilized to detect and analyze this phase separation behavior of this well-studied system. The distinction between the two phases, L_o_ and L_d_, is sensitive to the spatial localization of DPPC lipids and the number of DOPC neighbors. We designate DPPC and DOPC as primary and secondary lipids in the NMFk formalism, respectively. At the beginning of the simulations, there are no distinct phases and the lipid mixture is homogeneous. As simulation time progresses, lipids begin to segregate. By 0.6 us, the primary lipid type in this case, DPPC, begins to segregate into the L_o_ regions, while the secondary lipid type, DOPC, begin to segregate into the L_d_ domains. Thus, the number of secondary lipid neighbors around a given primary lipid should reflect that phase separation.

Given that a neighborhood profile of lipids can track the DOPC:DPPC:CHOL mixture, we first built a specific time-dependent data-matrix, ***X***_𝑳_(𝐭) (see Methods section) whose elements represent the number of DOPC neighbors to each one of the DPPC lipids in the system, within a specific radius, rcutoff extracted from careful analysis of the radial distribution function. NMFk decomposed the matrix ***X***_𝑳_(𝐭) into a product of two matrices: (i) the matrix of the basis lipid configurations, 𝑾_𝑵***K***_, whose columns present the configurations of DPPC lipids. NMFk determines the number of basis configuration ***K*** based on the robustness of the decomposition. Each one of the ***K*** basis lipid configurations in 𝑾_𝑵***K***_ contains the probabilities of the set of the DPPC lipids to have DOPC neighbors. The Silhouette-Reconstruction criterium was used to estimate the optimal number of basis lipid configurations to be 20. K-means clustering of each these 20 basis lipid configurations is demonstrating the tendency of increasing the number of primary lipids with a neglecting probability to have a neighbor of type DOPC when the time advances, which correspond to an increased total number of DPPC lipids located in nucleation domains.

By relating the basis lipid configurations to MD trajectories, we were able to show details of phase separation extracted by NMFk analysis. The NMFk discriminates lipids, depending on whether they belong to L_o_ or L_d_ phases or interfacial regions, as they undergo phase separation. Unlike, other analyses, basis lipid configurations provide details of lipids that take part in the nucleation versus those that establish line tension. It tracks the complicated features of the lipid segregation process leading to L_o_ and L_d_ phases. We identify lipids within the boundary of L_o_ phase with lipid configurations basis corresponding to a signature of L_o_ domain where DPPC is well packed and condensed by the high cholesterol concentration. Separately, another basis lipid configuration captures interfacial lipids that shield the DOPC lipids from such L_o_ domains during phase separation. Importantly, we demonstrate that the evolution of the nucleation process is captured in terms of lipid membership to different basis lipid configuration active in consecutive time intervals. NMFk identifies the lipids that take part in the initial nucleation and remains as part of the domain towards the phase separation. Also, other lipids that join such nucleation and remain till the phase separation are identified as the time progresses.

## CONCLUSIONS

The high variability and complexity of plasma membranes is still poorly understood. Higher resolution spectroscopy in combination with atomic detailed computer simulations are providing new insights, however, we are not yet close to fully understand or describe the membrane processes regulating cellular function. A detailed description of such complex systems undoubtedly requires novel mathematical frameworks that are able to decompose and categorize the evolution of thousands if not millions of lipids involved in the phenomenon.

Here we show the power of NMFk formalism on analyzing lipid phase separation and providing a robust analysis of categorizing lipids according to their localization in the membrane and elucidating the time-dependency along the nucleation process. The NMFk discriminates all different types of lipids, part of L_o_ or L_d_ or interface, due to their particular behavior along the trajectory and the resulting probability to have a DOPC neighbor. If there is no clear pattern in the behavior of the lipids, for example, when MD simulations do not produce any distinct behavior associated with phase separation, NMFk analysis does not produce false features. This is the first demonstration of NMFk serving as a useful tool in detecting time-dependent domain formation and lipid separation in MD simulations of complex lipid mixture systems. Even though we have exhibited the usefulness of NMFk in the context of a well-studied ternary lipid mixture, an extension to more complicated lipid mixtures is feasible with a tensor formalism and is currently under consideration.

## MATERIALS AND METHODS

### Membrane patch

An initial configuration of a CG membrane patch was obtained using the script tools provided in the Martini force field website (http://cgmartini.nl/). Our CG lipid system contains DPPC:DOPC:CHOL lipids in a 37:36:27 ratio respectively, which initially were randomly placed within a XY plane. This lipid ratio has been experimentally and computationally observed to transitioning towards a phase separated Liquid-ordered/Liquid-disordered state ^54, 56^. The lipids were represented using the Martini V2.2 force field ^39^ with a refined set of parameters, which has shown an improved phase separation behavior ^52^. Similarly, simulation with the standard Martini lipid model was also carried on. The total system was composed of 16366 lipids, 718830 Martini water beads (175 atomistic water molecules per lipid) and 150mM NaCl to preserve an overall constant ionic strength. In order to avoid spontaneous freezing of the Martini water beads (a well-known artifact previously reported in the original model ^39^), 0.1% M of the water beads were replaced by anti-freeze particles.

### Molecular dynamics protocol

We followed a current update in parameters set-up for performing the CG simulations ^57^. The equations of motion were integrated every 30fs time-step. A reaction-field electrostatics algorithm was used with a Coulomb cut-off of 1.1 nm and dielectric constants of 15 or 0 within or beyond this cut-off, respectively. Lennard-Jones interactions were cut off at 1.1 nm, where the potential was shifted to zero. In order to accelerate the lipid phase de-mixing, constant temperature was maintained at 290 K via separate coupling of the solvent (water and ions) and membrane components using a velocity-rescaling thermostat ^58^ with relaxation times of 1.0ps. During equilibration, the Martini beads representing the phosphate groups of the lipid head regions were positionally (xyz components) restraint in order to preserve the initial random positions. In this stage, the solvent molecules (water and ions) were allowed to diffuse and the box pressure was maintained semi-isotropically coupled at 1 bar using the Berendsen barostat ^59^ with relaxation times of 12 ps and compressibilities of 3 × 10^−4^ bar^−1^. After that, production runs were performed using a Parrinello-Rahman barostats ^60^. Simulations were run for 20 μs using the GROMACS version 5.2.1 ^61^ and the trajectories were saved every 3 ns providing the frames for the construction of the NMFk matrix (see later).

### Generation of the contact matrix 𝐗_𝐍_(𝐭)

Every frame stored within 2 us (667 frames in total) were used for generating the corresponding matrix for NMFk analysis. We rely on the implemented GROMACS tool *gmx select* to output the number of DOPC lipids around every DPPC molecule within 1.1 nm. This cutoff-radius structurally corresponds to the second layer of neighbors, as estimated by the second maximum peak of the radial distribution function ***g***(r) (**Fig. 3**). Thus, each column of the data-matrix, 𝑋_*N*_(𝐭), corresponds to a variable number of the DOPC neighbors of a given DPPC lipid per frame, while the rows correspond to the number of the 3038 DPPC lipids in the system. Similarly, matrix reconstruction was carried out for 20 us collection. An example of the matrix can be found as part of Supporting material.

### Nonnegative Matrix Factorization and NMFk

Nonnegative Matrix Factorization (NMF) is a well-known unsupervised machine learning method created for parts-based representation ^36^. NMF has been successfully leveraged for decomposing mixtures of various types nonnegative signals, i.e., for Blind Source Separation (BSS) ^38^ problems. If the BSS problem is solved in a temporally discretized framework, the goal of the NMF algorithm is to retrieve the original nonnegative signals (sources), 𝑾; 𝑾 ∈ *M*_𝑃𝐾_(*R*_+_) that produced the observational records, ***X***; ***X*** ∈ *M*_*N*𝑃_(*R*_+_), detected at a set of sensors. Here 𝑅+ denotes the set of real nonnegative numbers, *N* is the number of the recording sensors, ***K*** is the number of unknown original signals, and 𝑃 is the number of discretized moments in time (time points or frames) at which the signals are recorded at the sensors. Only the matrix ***X*** is known initially. Thus, in a BSS problem, the recorded data, ***X***, is formed by a linear mixing of ***K*** unknown original signals 𝑊, blended by an unknown mixing matrix, 𝑯. Since both factor matrices 𝑾 and 𝑯 are unknown, and even their size ***K*** (i.e., the number of unknown original signals) is unknown the problem is typically under-determined. NMF can solve such kind of problems by leveraging, for example, the multiplicative update algorithm ^37^ to minimize the Frobenius 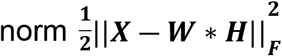. An additional advantage of NMF method is that it can work with data in which the original signals are not independent but partially correlated ^38^.

One of the difficulties of the NMF algorithm is that it requires prior knowledge of ***K***- the number of the unknown original signals. Recently a new protocol called NMFk addressing this limitation has been reported ^62-64^. This protocol complements classical NMF with custom k-means clustering and Silhouette ^65^ statistics, which allows simultaneous identification of the optimal number of the unknown basis patterns. The NMFk was utilized to successfully decompose the largest available dataset of human cancer ^66^ genomes, as well as for extraction of physical pressure transients ^64^ and contaminants ^67^ originating from an unknown number of sources that may propagate with a finite speed in nondispersive ^68^ or dispersive media ^69^ as well as for extraction of the original crystal structures and phase diagram from X-ray spectra of material combinatorial libraries ^70^.

NMFk determines the number of the unknown original signals based on the robustness and reproducibility of the NMF solution. Specifically, it explores consecutively the possible numbers of configurations ***K̃*** (***K̃*** can go from 1 to N-1, where N is the total number of frames), by obtaining sets of NMF minimization solutions for each ***K̃***. Note that ***K*** serves to index the different NMF models, and it is distinct from ***K***, which is fixed, albeit unknown number. Further, NMFk leverages a custom clustering using the cosine similarity, in order to estimate the robustness of each set of NMF solutions with fixed 𝐾 but derived with different initial guesses. Comparing the quality of the derived clusters (a measure how different are the extracted signals) and the accuracy of minimization among the sets with various ***K̃***, which we call a Silhouette-Reconstruction criterium, NMFk determines the optimal numbers of the unknown original signals. To access the quality of the clusters obtained for each set we use their average Silhouette width, 𝑺. NMFk utilizes 𝑺 to measure how good is a particular choice of ***K̃*** as an estimate for ***K***. Specifically, the optimal number of patterns is picked by selecting the value of ***K*** that leads to both: (a) an acceptable reconstruction error ***R*** of the observation matrix ***X***, where

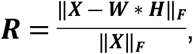

and (b) a large average Silhouette width (i.e., an average Silhouette width close to one). The combination of these two criteria is easy to understand intuitively. For solutions with ***K̃*** less than the actual number of patterns (***K*** < ***K̃***) we expect the clustering to be good (with an average Silhouette width close to 1), because several of the actual patterns could be combined to produce one “super-cluster”; however, the reconstruction error will be high, due to the model being too constrained (with too few degrees of freedom), and thus on the under-fitting side. In the opposite limit of over-fitting, when ***K̃* > *K*** (***K̃*** exceeds the actual number of configurations), the average reconstruction error could be quite small - each solution reconstructs the observation matrix very well - but the solutions will not be well-clustered (e.g., with an average Silhouette substantially less than 0.8), since there is no unique way to reconstruct ***X*** with more than the actual number of configurations, and no well-separated clusters will be formed.

Thus, the best estimate for the number of unknown original signals, 𝑾, corresponding to the true number of unknown original signals ***K***, is given by the value of ***K̃*** that optimizes both of these metrics simultaneously. Finally, after determining ***K***, we use the centroids of the ***K*** clusters as a final robust representation of the original signals.

### NMFk minimization algorithm

Here we leveraged the multiplicative algorithm ^37^ based on Kullback–Leibler divergence ^71^ as well as the block coordinate descent algorithm ^72^ based on Frobenius norm. We did not observe any significant difference between the results obtained via these two algorithms.

### NMFk clustering algorithm

NMFk creates up to N-1 sets of minimizations (called NMF runs), one for each possible number ***K̃*** of original patterns. In each of these runs, 𝑸 solutions (e.g., 𝑸=200) of the NMF minimizations for a fixed number of patterns ***K̃*** are derived. Thus, each run results in a set of solutions *U*_*K̃*_:

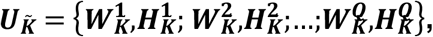

where each of these “tuples” represents a distinct solution for the nominally same NMF minimization: the difference is stemming from the different (random) initial guesses. Next, NMFk performs custom clustering, assigning the ***K̃*** columns/features of each **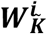** of all 𝑸 solutions to one of the ***K̃*** clusters, representing ***K̃*** basic patterns. This custom clustering is similar to k-means clustering but with an additional constraint which holds the number of elements in each of the clusters equal to the number of solutions 𝑸. For example, with 𝑸=200 each one of the ***K̃*** identified clusters must contain exactly 200 solutions. This condition has to be enforced since each minimization (specified by a given **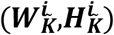** tuple) contributes only one solution for each feature, and accordingly supplies exactly one element to each cluster. During the clustering, the similarity between patterns is measured using the cosine similarity.

### Numerical codes, data and Supporting Information

The following files: COORD.gro with the coordinates; INPUT.mdp with the parameters set to start the simulation with Gromacs; PARAMETERS.top with the Martini force field-based topology for the membrane system; DPPC-DOPC-1.1nm.xvg with an example matrix generated from the MD trajectories used for NMFk calculation, are also provided as supporting information accompanied this paper.

To extract patterns of the basis lipid configurations from the MD simulations we used the NMFk method which is an extension of the original NMF ^37^ that includes a custom clustering for determination of the number of patterns ^66^. Our NMFk protocol is based on the SigProfile software created for identification of mutational signatures in human cancer66.

The SigProfile code is freely available at: https://www.mathworks.com/matlabcentral. To use SigProfile, an input file should be at place. In our case, the input file is the contact matrix ***X***_𝑵_(𝐭) with size (***N*** × ***M***), where ***N*** is the number of the lipids in the MD simulations and ***M*** is number of the frames. A detailed description of NMFk is available elsewhere ^64^. The input data-file, containing the contact matrix ***X***_𝑵_(𝐭) as well as a script, needed to run the SigProfile, are provided as supplemented information accompanied this paper. A README.docx file with a list of included files, links to publicly available repositories along with their brief description and instructions is also provided as supporting information.

The simulated MD-data containing the lipids’ trajectories (∼ 100GB) is available freely but because of its size-upon request to.

## ACKNOWLEDGMENTS

This research was performed at Los Alamos National Laboratory and carried out under the auspices of the National Nuclear Security Administration of the United States Department of Energy. We like to acknowledge Tyler Reddy for generating synthetic data to test preliminary results. The work was supported by LANL LDRD grant 20180060DR. This work has been supported in part by the Joint Design of Advanced Computing Solutions for Cancer (JDACS4C) program established by the U.S. Department of Energy (DOE) and the National Cancer Institute (NCI) of the National Institutes of Health. This work was performed under the auspices of the U.S. Department of Energy by Lawrence Livermore National Laboratory under Contract DE-AC52-07NA27344, Los Alamos National Laboratory under Contract DE-AC5206NA25396, Oak Ridge National Laboratory under Contract DE-AC05-00OR22725, and Frederick National Laboratory for Cancer Research under Contract HHSN261200800001E. We thank the LANL Institutional Computing for the computing resources.

## CONTRIBUTIONS

All authors designed the research. C. A. L. performed the MD simulations. B. S. A. performed the NMFk analyses. All authors analyzed the data and wrote the manuscript.

